# A Minimum Perturbation Theory of Deep Perceptual Learning

**DOI:** 10.1101/2021.10.05.463260

**Authors:** Haozhe Shan, Haim Sompolinsky

**Author notes:** http://hzshan.github.io.

## Abstract

Perceptual learning (PL) involves long-lasting improvement in perceptual tasks following extensive training and is accompanied by modified neuronal responses in sensory cortical areas in the brain. Understanding the dynamics of PL and the resultant synaptic changes is important for causally connecting PL to the observed neural plasticity. This is theoretically challenging because learning-related changes are distributed across many stages of the sensory hierarchy. In this work, we modeled the sensory hierarchy as a deep nonlinear neural network and studied PL of fine discrimination, a common and well-studied paradigm of PL. Using tools from statistical physics, we developed a mean-field theory of the network in the limit of large number of neurons and large number of examples. Our theory suggests that, in this “thermodynamic” limit, the input-output function of the network can be exactly mapped to that of a deep *linear* network, allowing us to characterize the space of solutions for the task. Surprisingly, we found that modifications of synaptic weights in lower levels of the hierarchy are both sufficient and necessary for PL. To address the degeneracy of the space of solutions, we postulate that PL dynamics are constrained by a normative “minimum perturbation” (MP) principle, which favors weight matrices with minimal changes relative to their pre-learning values. Interestingly, MP plasticity induces changes to weights and neural representations in all layers of the network, except for the readout weight vector. In addition, such plasticity can be learned simply through slow learning. We further elucidate the properties of MP changes and compare them against experimental findings. Overall, our statistical mechanics theory of PL provides mechanistic and normative understanding of several important empirical findings of PL.

## I. INTRODUCTION

Perceptual learning, the improvement of performance in perceptual tasks after practice, is one of the most basic forms of learning in the brain and has been extensively studied experimentally [1–8]. Physiologically, PL is accompanied by long-lasting changes to neuronal response properties in cortical areas. Connecting physiological changes to behavioral observations has been challenging, in part due to the complex learning dynamics and processing in the sensory hierarchy, which is composed of multiple cortical regions. As a result, several important issues concerning the neural mechanisms of PL remain unresolved after decades of research.

First, which cortical areas undergo modifications and which of the changes causally drive PL? While behavioral specificity of PL [9, 10] points to an important role for plasticity in early sensory areas, single-unit response properties in visual areas V1 and V2 show only minor changes after PL [1, 2]. In addition, PL induces significant changes to single-neuron properties in intermediate to late stages of visual processing, such as V4 [3, 4, 7, 8], LIP [11], and IT [12, 13].

Second, what are the functional consequences of the observed changes? Analysis of changes in neuronal responses after PL indicates improved accuracy of the neural coding of the trained stimuli [6–8]. This appears to be inconsistent with the behavioral finding that PL does not transfer to a different task even when using the same stimuli [14–16]. The Reverse Hierarchy Theory [17] proposes that PL is initially driven by learning in high areas, which results in less specific learning; modifications of lower areas follow if the task is difficult, as for instance in fine perceptual discrimination tasks, leading to more specific learning. Analysis of a reduced model of perceptual learning has lent support for this theory [18]. However, recent experimental and computational studies questioned these predictions [6, 7, 19], providing evidence of changes in primary sensory areas already in the early stages of PL. On the other hand, experiments in random dot visual motion discrimination tasks found that PL is correlated with changes in decision-making areas (e.g., LIP) but not sensory areas (e.g., MT) [11, 20]. From a theoretical perspective, the hierarchical nature of the underlying sensory system implies that there is an enormous degeneracy of possible synaptic weight matrices that solve the task of PL.

Most existing theories of PL assume changes only in either the weights of the readout from a fixed sensory array [21–23] or the input layer to a single cortical circuit [19]. Such “shallow” models are inconsistent with the sensory hierarchy in the brain and do not address the neural correlates of PL in multiple cortical regions.

In the present work, we directly addressed the issue of PL in a deep network by studying PL of a fine discrimination task in a deep neural network (DNN) model of the sensory hierarchy [24–26]. As learning dynamics in DNNs are in general challenging to study [27–35], we developed a mean-field theory of information propagation in the model at the limit of large numbers of neurons in every layer and large number of training examples. The theory reveals that during the perceptual task, the DNN effectively behaves like a deep *linear* neural network. This considerably simplifies the theoretical analysis of the space of solutions, as well as the emergent changes in neural representations. Surprisingly, We found that modifications of synaptic weights in the first level of the hierarchy are both sufficient and necessary for PL. To address the degeneracy of the space of solutions, we developed a *normative* theory of PL. Specifically, we postulated that in the brain, learning dynamics are constrained by a normative “minimum perturbation (MP)” principle, which favors weight matrices with minimal changes relative to their pre-learning values. Interestingly, MP learning induces changes in weights and neural representations in all layers of the networks, except for the readout weight vector. MP learning predicts changes to tuning properties of cortical neurons that are consistent with experimental observations and suggests that signal amplification, not noise reduction, is the primary driver of PL. Our theory makes the readily testable prediction that PL can simultaneously lead to positive and negative transfer to different untrained stimuli. Finally, we found that MP learning can be implemented through slow gradient-descent learning. Overall, leveraging the large size of the network involved in PL, we have developed a statistical mechanics theory of PL in deep neural networks which provides mechanistic and normative understanding of several important empirical findings of PL. This work complements recent theoretical studies of learning in deep networks [27–35], contributing to the understanding of learning and computation in these important architectures.

Our deep network model of PL is described in Section II. The mean field analysis is summarized in Section III. Section IV presents the MP principle and analyzes PL with minimum perturbation. Section V analyzes the use of gradient descent to learn MP plasticity. A discussion of the implications for the field of perceptual learning is presented in Section VI.

## II. A DEEP NETWORK MODEL OF PL

### A. Input channels

We assume *N* input channels (**Fig. 1A**, gray squares) representing a 1D stimulus. Neurons in the input channels are indexed by a preferred stimulus angle 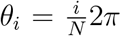 for the *i*th neuron. The collective response of input channels to a stimulus with angle *θ* is given by the *N*-dim vector

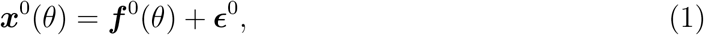

where ***ϵ***^0^ is i.i.d. Gaussian noise with zero mean and variance *σ*^2^. The noise averaged response of each input neuron is given by a bell-shaped tuning curve centered on its preferred stimulus

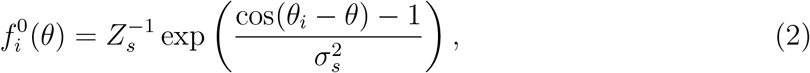

where *Z_s_* ensures 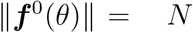 and *σ_s_* controls the input selectivity, assumed to be the same for all channels (**Fig. 1C**). Tuning and noise properties of input channels are not affected by learning.

**FIG. 1.**
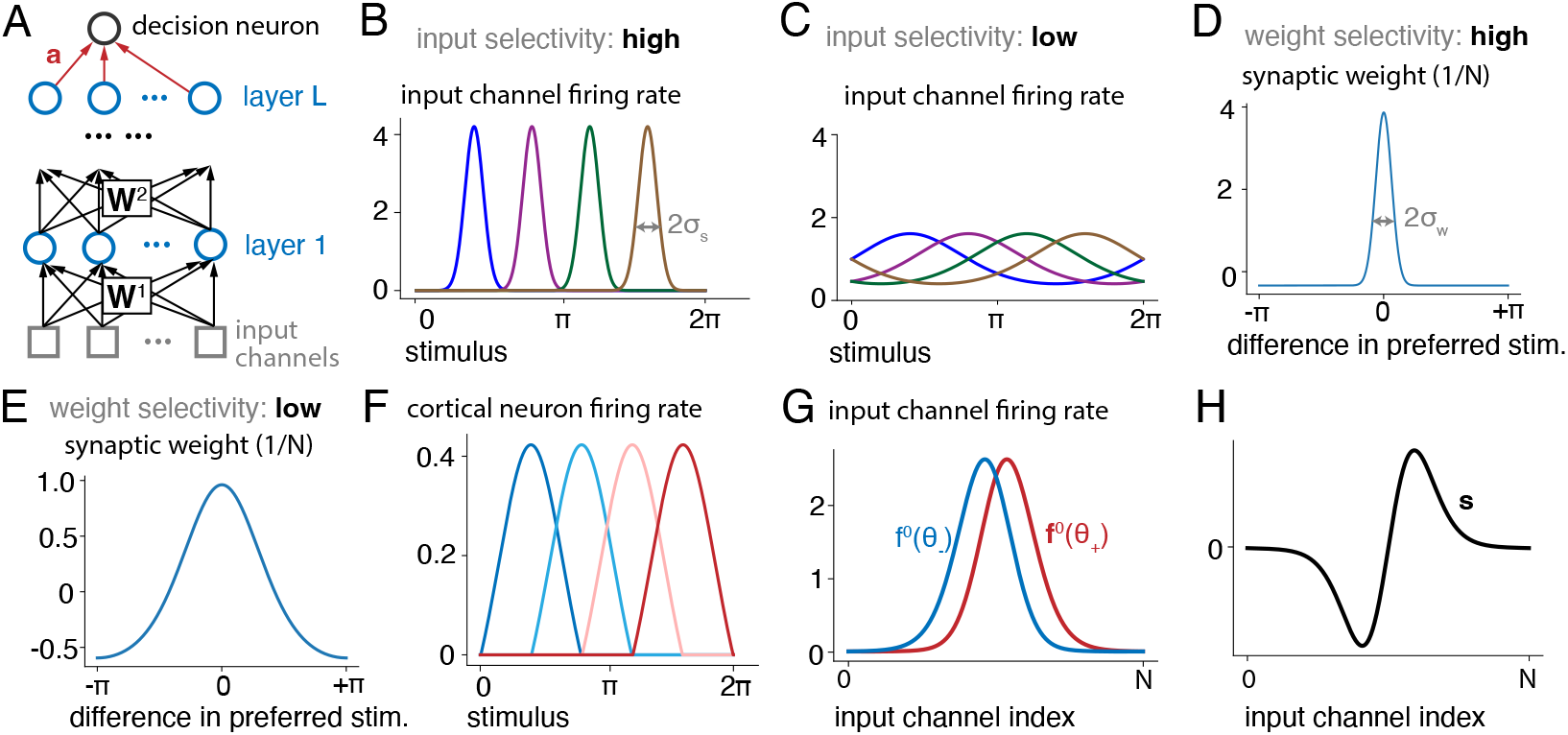
Model of perceptual learning. **A** Diagram of the deep network model of the sensory hierarchy. The output of input channels (gray squares) is passed through *L* layers of cortical neurons (blue circles) before getting read out by a linear readout (***a***). **B** Example tuning curves of input channels. Each curve represents a channel with a different preferred stimulus. The preferred stimuli of input channels uniformly tile [0, 2*π*]. This panel shows the regime of high input selectivity and hence narrow tuning curves. **C** Same as B, but for the scenario of low input selectivity. **D** Example feedforward weight structure before learning. Weights connecting neurons with similar preferred stimuli tend to be excitatory (positive) and strong while those connecting neurons with dissimilar preferred stimuli tend to be weak and inhibitory (negative). This panel shows the regime of high weight selectivity. **E** Same as D, but for low weight selectivity. **F** Bell-shaped tuning curves of input channels and the initial weight patterns lead to bell-shaped tuning curves for all cortical neurons before PL. **G** Noise-averaged activity of the input channels (***f***^0^(*θ*)) in response to the two presented stimuli, *θ*_±_. The difference between them is exaggerated here for illustration purposes. **H** The signal ***s*** is in the direction of the difference between ***f***^0^(*θ*_±_).

### B. Model architecture and pre-PL weights

Our model of the sensory system is a feedforward network with *L* hidden layers and a linear readout from the top layer (**Fig. 1A**). Each hidden layer is composed of *N* rectified linear (ReLU) neurons (“cortical neurons”). Let ***x**^l^*(*θ*) denote the noisy population response vector of neurons in layer *l*, and ***f**^l^*(*θ*) its average over noise. {***x**^l^*(*θ*)}_*l*=1,.,…,*L*_ are recursively given by

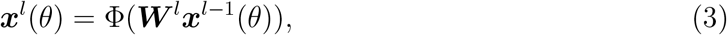

where Φ(·) is the element-wise rectified linear function. The linear behavioral readout ***a*** produces a scalar network output from activity in the last layer

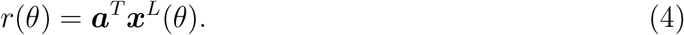

Pre-PL weights are modelled after feedforward synaptic connections between visual areas in the brain. We chose a circulant structure which is appropriate for propagating angular signals. Every neuron receives strong, excitatory input from neurons in the previous layer with similar preferred stimuli and weak, inhibitory input from neurons with dissimilar preferred stimuli. Concretely, pre-PL weights {***W**^l^*}_*l*=1,2,…,*L*_ are given by

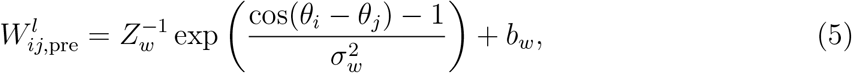

where *Z_w_* is chosen such that each row of 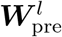 has norm 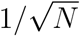 (i.e. each weight is *O*(*N*^−1^)). This normalization ensures that the input to any hidden neuron is of magnitude *O*(1). The offset *b_w_* is chosen such that each row sums to 0. The parameter *σ_w_* controls selectivity of the pre-learning weights; small *σ_w_* leads to a high-selectivity weight structure where a few weights dominate the input (**Fig. 1D**) and vice versa (**Fig. 1E**). As a result of the input tuning curves and the feedforward weight structure, all cortical neurons are tuned to the 1D stimulus and have bell-shaped tuning curves before learning (**Fig. 1F**).

### C. Fine-discrimination task

We focus on learning a fine discrimination task, where one out of two similar visual stimuli is presented to the subject, who must correctly indicate which one is presented. In our model, the task consists of discriminating two values of the stimulus, *θ*_±_ = *θ*_tr_ ± *δθ*, where the center stimulus *θ*_tr_ is called the trained stimulus and *δθ* ~ *O*(*N*^−1/2^). This choice of scaling ensures that the total signal-to-noise ratio (SNR) in the input layer is *O*(1). In each trial, one of *θ*_±_ is presented and generates a noisy activation of the input array (Eq. 1, **Fig. 1G**). In each trial, the decision neuron activity *r* indicates whether the input comes from the *θ*_+_ stimulus or from *θ*_−_ with *r* > 0 or *r* < 0, respectively. Stimuli are presented with equal probability; the optimal performance in the task is thus given by performing maximum likelihood discrimination (MLD [21]). Importantly, since the noise is Gaussian, the task can be performed optimally by a linear discriminator reading out directly from the input channels and using weights parallel to the signal (**Fig. 1H**), defined as the unit vector

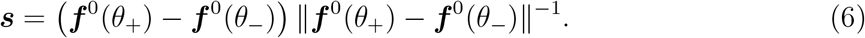

Thus the output in this scenario equals ***s**^T^**x***^0^(*θ*) which leads to optimal performance in this setup [36].

### D. Pre-PL readout

We assume the pre-PL value of the readout weight vector ***a***_pre_ to be optimized for this task when reading out the pre-PL top-layer representations. Thus, we initialize the pre-PL readout such that it minimizes the loss function between the network readout and the optimal output (see below and S.M. I). The rationale for non-random initialization of the readout weights is to provide the network with well-above-chance but generally suboptimal performance (as shown below, it is suboptimal because the top-layer representations may be suboptimal). In the context of animal experiments, this mimics the situation where animals understand the task but have not yet acquired the expert skills required for near optimal performance.

### E. Learning

We model the process of PL as modifying weights in order to minimize the discrimination error. Since the optimal output for this task is given by ***s**^T^**x***^0^(*θ*) it is convenient to use a mean-squared error objective function

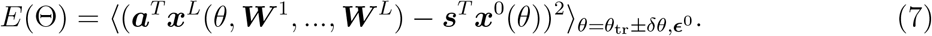

where Θ = (***W***^1^, ***W***^2^, …, ***W**^L^, **a***) denotes the vector of all weights of the networks and angular brackets denote averaging over the two stimuli and noise. This cost function measures the deviation of the input-output function of the system from the optimal one.

## III. A MEAN-FIELD THEORY OF PL OF THE FINE DISCRIMINATION TASK

In this section, we describe our mean-field approach to studying sensory processing and PL in the deep, nonlinear network by approximating it with an equivalent linear network. We describe the approximation (Sec. III A) and the insights it provides into why pre-PL cortical representations can be suboptimal (Sec. III B). We discuss the space of possible solutions that this theory revealed in Sec. III C.

### A. Equivalent linear networks

First, we note that, during the fine discrimination task, signal and noise-induced fluctuations in the input to any neuron are small (they both scale as *N*^−1/2^). This can be seen by considering the scaling relations *δθ* ~ *O*(*N*^−1/2^), *σ*^2^ ~ *O*(1) and 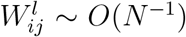. At the large *N* limit, we can expand activities of cortical neurons around their average inputs by writing (using ⊙ to denote the Hadamard product and ***f**^l^* to denote ***f**^l^*(*θ*_tr_); this is similar to the approximation done in [37] for recurrent networks)

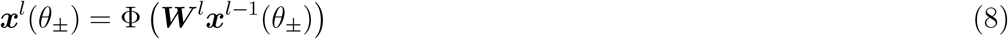

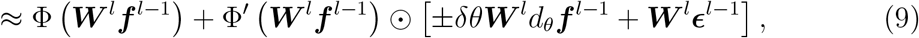

where

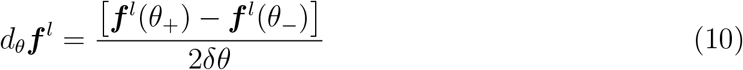

and 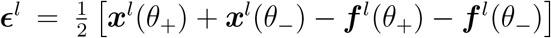 are the signal-induced and noise-induced fluctuations in layer *l*, respectively. At large *N*, by the centra limit theorem, the components of noise are Gaussian (though correlated). For ReLU nonlinearity, the activation slope 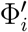 is 1 for an *active* neuron and zero for an *inactive* one. At the limit of large *N*, fluctuations in the input to each neuron are small compared to the mean. Thus, inactive neurons remain quiescent for most of the trials and do not contribute to the network output. For a similar reason, activities of active neurons are [***W**^l^**f***^*l*−1^]_*i*_ i.e, they are *linear* functions of activities of neurons in the previous layer. Thus, we can define effective weight matrices, 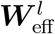, as

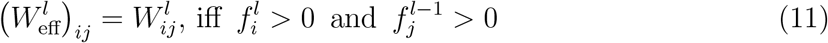

and zero otherwise. Importantly, this approximation holds only during fine discrimination around a fixed *θ*_tr_ because of the strong similarity between the different inputs. Inputs with angles very different from *θ*_tr_ will be processed by different sets of effective weights. Given the analysis above, during the fine discrimination task around *θ*_tr_ the input output function of the deep network is effectively linear,

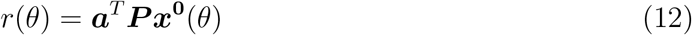

where

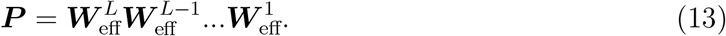

We call *P* the *processing matrix* (**Fig. 2A**, right). We proceed to consider how the properties of *P* affect task performance.

**FIG. 2.**
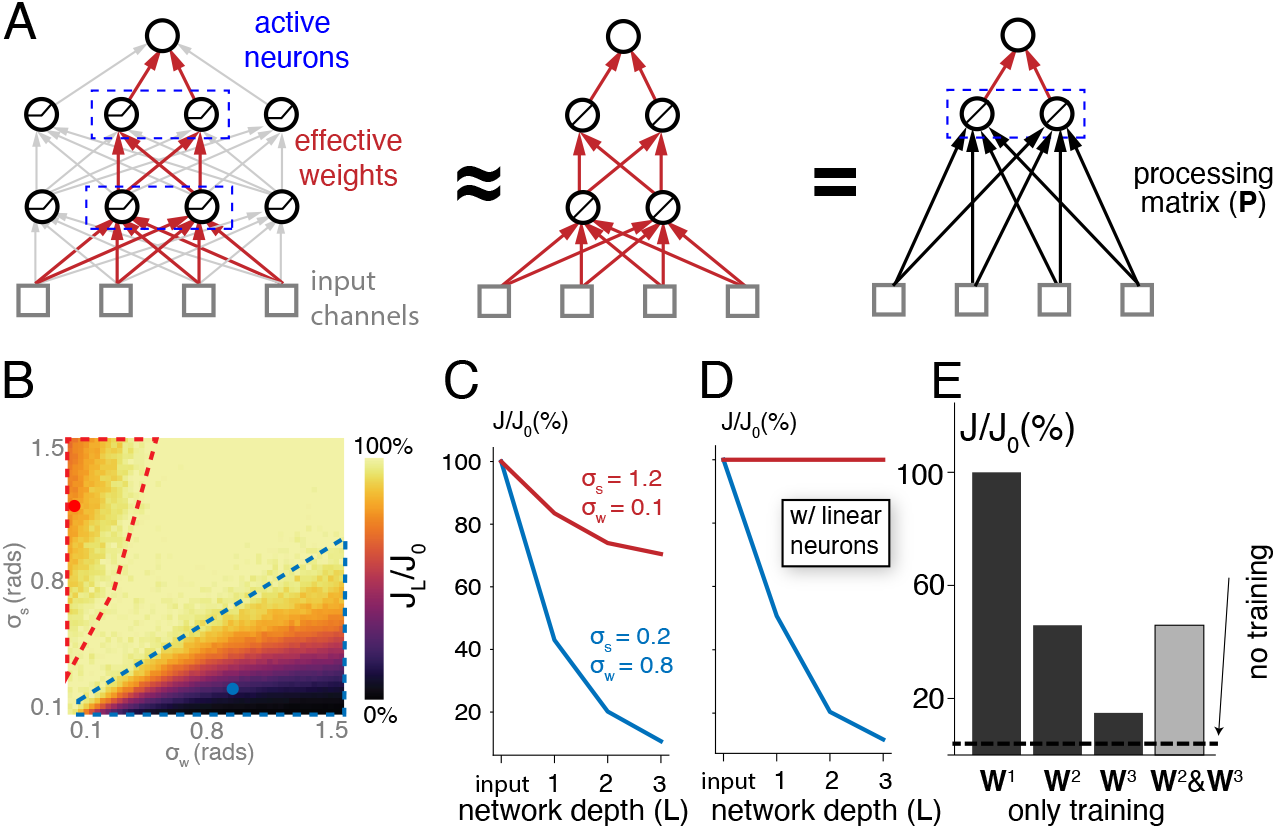
Suboptimal neural representations before learning. **A** Schematics showing the relationship between weights (all arrows, left), effective weights (red arrows, left and center), and the processing matrix (black arrows, right). **B** Information for the trained stimulus in the last layer (*J_L_*) divided by the input information (*J*_0_), for different input and weight selectivity. The ratio is low for large *σ_s_*, small *σ_w_*(red polygon) or small *σ_s_*, large *σ_w_* (blue polygon). Dots: example parameters used in C, D. *N* = 1000 in all panels. *L* = 1 in this panel. See Fig. S1 for deeper networks. **C** Information for the trained stimulus in the last layer of networks of different depths, divided by the input-layer information (*J*_0_). **D** Same as E, but assuming that all neurons are active. **E** Best last-layer information achievable if plasticity is restricted to some weight matrices in a three-layer network. Dashed line: performance if no weight matrix is modified. Modifying any weight matrix improves the performance, but only modifying ***W***^1^ is sufficient and necessary for optimizing it.

### B. Pre-learning suboptimal representations

Optimizing ***a***_pre_ amounts to optimizing a linear readout from an input ***Px***^**0**^(*θ*) which contains a signal and an additive (correlated) noise. In such a system, the probability of error under optimal readout is given by 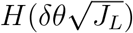 where 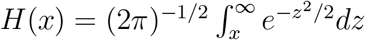 and *J_L_* is the linear Fisher information [38]. It is defined as

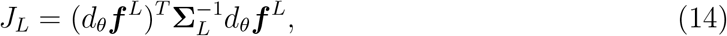

where the matrix **Σ**_*L*_ is the noise covariance matrix in the top layer. Given the linear approximation, it is given by **Σ**_*L*_ = *σ*^2^***PP**^T^* and the top layer signal is *d_θ_**f**^L^* = ***P**d_θ_**f***^0^. Even if ***P*** is low rank (see below), the (pseudo)-inverse (***PP**^T^*)^−1^ is well defined when multiplied by ***P***. Hence,

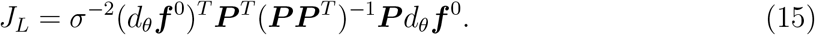

Note that ***P**^T^* (***PP**^T^*)^−1^***P*** is a projection matrix. It is identity if ***P*** is full rank. Otherwise, it projects inputs onto the low-rank subspace spanned by its rows. Thus, Eq. 15 states that *σ*^2^*J_L_* is the squared norm of the projection of the signal vector *d_θ_**f***^0^ ∝ ***s*** onto the subspace spanned by ***P***. The network is optimal if ***s*** resides in the span of ***P***, yielding *J_L_* = *J*_0_ = *σ*^−2^||*d_θ_**f***^0^||^2^.

We now ask whether the pre-PL weights are already optimal for the present task. We computed the singular value decomposition of the pre-PL ***P*** and found that it has a low rank structure (Fig. S2). There are two sources of the reduced rank of ***P*** depending on the system parameter regime: a “selective-input-unselective-weights” regime characterized by small *σ_s_* and large *σ_w_* (**Fig. 2B**, blue) and an “unselective-input-selective-weights” regime, characterized by large *σ_s_* and small *σ_w_* (**Fig. 2B**, red). In the selective-input-unselective-weights regime, the pre-PL network weights 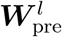 are low rank (even before rectification) due to the smoothness of circulant weights, implying that they project to subsequent layers only part of the signal in the input. In this regime, information loss occurs regardless of the rectification of representation neurons (**Fig. 2C, D**, blue line). On the other hand, in the unselective-input-selective-weights (large *σ_s_*, small *σ_w_*) regime, the original weight matrices project the full signal. However, due to firing-rate rectification, a substantial fraction of the neurons are inactive for essentially all training stimuli. Thus, the effective weights are low rank. In this regime, the low-rank structure of ***P*** disappears if we remove neuronal rectification (**Fig. 2C, D**, red line). In both cases, the signal contains a substantial component perpendicular to the low-rank span of pre-PL ***P***, as evidenced by computing *J_L_*/*J*_0_ (**Fig. 2B**). Hence the pre-PL network exhibits suboptimal performance.

### C. Space of solutions

We derived the following *necessary and sufficient condition* on post-learning effective weights that accomplish this goal (derivations in S.M. II; hereafter we use ***W**^l^* to denote 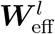 for brevity): for any values of ***W***^2^, …, ***W**^L^* and ***a*** that satisfy ***ã*** ≡ (***W***^2^)^*T*^ …(***W**^L^*)^*T*^ ***a*** ≠ 0, the task can be performed optimally if and only if ***W***^1^ satisfies

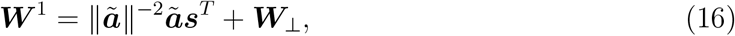

where 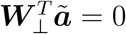. This result implies that the learning problem can be solved for essentially arbitrary (non-zero) higher-layer weights as long as ***W***^1^ is adjusted accordingly. Conversely, restricting the plasticity to higher-layer weights while freezing the first layer weights to their pre-PL values does not obey this condition; thus this is insufficient for optimal performance, as discussed earlier (**Fig. 2E**). This result underscores the critical role played by early sensory areas in PL.

## IV. LEARNING WHILE MINIMIZING NETWORK PERTURBATION

The large space of solutions makes it hard to predict the pattern of changes in the circuit induced by learning. To remove this degeneracy, we propose an optimality criterion, “minimum perturbation” (MP), that favors a solution with small perturbations to pre-PL weights. According to this criterion, the optimal weights are

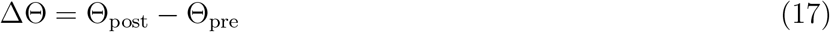

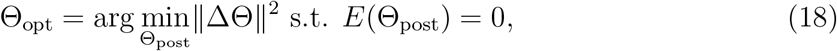

where “post” indicates post-PL weights. This principle is motivated by the brain’s need to maintain relatively stable representations while learning a new task. We analytically solved this optimization problem with Lagrange multipliers by first setting up the Lagrangian

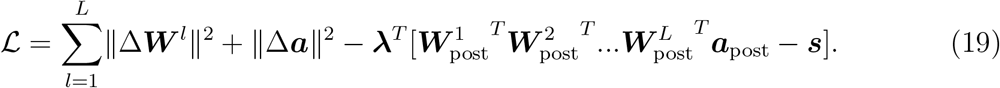

Extremizing the Lagrangian w.r.t. weight changes reveals a general rank-1 structure for MP Δ***W**^l^*

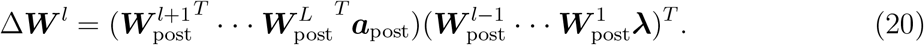

Solving the above equations requires introducing 2*L* scalar order parameters, which obey 2*L* self-consistent equations that need to be solved numerically. Expressions of the order parameters and self-consistent equations for *L* = 1, 2, 3, as well as the numerical procedure for solving the self-consistent equations, are given in S.M. III.

We discuss features of the solutions below.

### Distribution of MP plasticity

We quantified the magnitude of MP modification to weights in each layer by computing 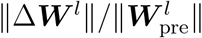 (**Fig. 3A**; see Fig. S5 for the unselective-input-selective-weights regime). The analysis reveals two salient features that are consistent across system parameters. First, MP plasticity predominantly affects lower-layer weights. Second, surprisingly, MP plasticity does not appreciably alter the readout (**Fig. 3A**, red line). This suggests that hidden-layer representation changes, rather than readout changes, drive performance improvement. Plasticity in higher-layer weights ***W***^*l*≥2^ plays the important role of reducing the overall perturbation to the network. Indeed, if we restrict learning to ***W^1^***, the total perturbation is greater (**Fig. 3B**).

**FIG. 3.**
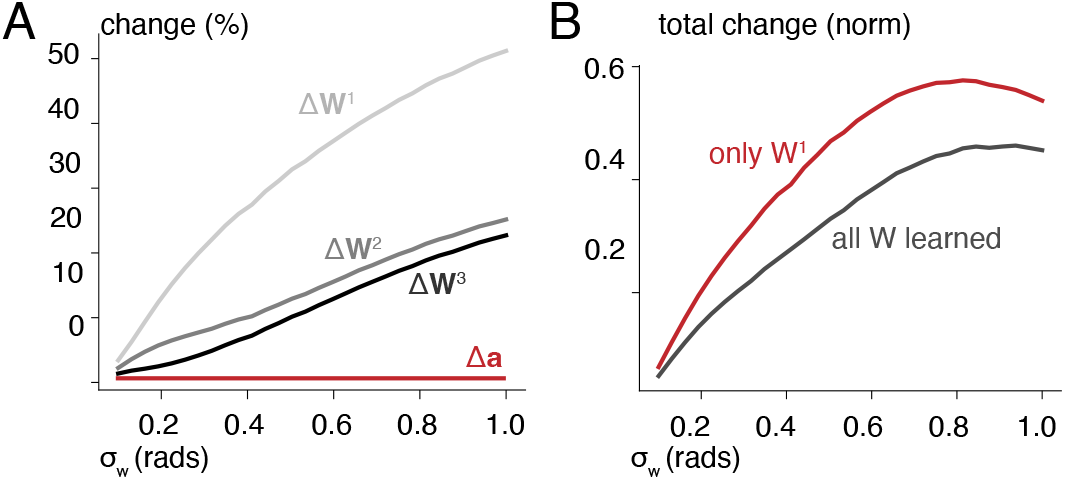
Minimum-perturbation plasticity of perceptual learning. **A** The magnitude of synaptic changes to each matrix and the readout vector ***a***, for networks initialized with different *σ_w_*. Percent change is defined as the Frobenius norm of synaptic changes divided by that of the pre-PL weight matrix. In all panels, *σ_s_* = 0.2, *L* = 3, *N* = 1000. **B** Restricting learning to ***W***^1^ leads to more network-wide perturbation (measured by the sum of matrix norms of Δ***W***^1^, Δ***W***^2^, Δ***W***^3^) than unrestricted learning. In either case, the readout ***a*** is also allowed to learn but does not change significantly following PL.

### Performance improvement is driven by signal amplification

An important and long-standing debate in PL research is whether behavioral improvement is driven by signal amplification, noise suppression, or both [1, 3, 4, 7, 39–42]. To address this question within the framework of MP learning, we define the signal and noise contributions via *J_L_* = (signal/noise)^2^, where the signal amplitude is ||*d_θ_**f**^L^*|| and the noise amplitude is defined via noise^−2^ = (*d_θ_**f**^L^*)^*T*^**Σ**_*L*−1_*d_θ_**f**^L^*/||*d_θ_**f**^L^*||^2^. The network after MP learning exhibits a pronounced amplification of signal (**Fig. 4A**), with the effect being stronger in higher layers. Surprisingly, we found that PL also *amplifies* noise across all layers, although to a weaker extent than signal amplification (**Fig. 4B**; effects on noise correlation are shown in Fig. S4). Thus, MP learning improves perceptual performance by strengthening the signal rather than weakening the noise. This analysis also reveals that signal/noise changes are generally greater in higher layers even though weight changes are greater in lower layers, highlighting the difference between distribution of weight changes and distribution of representation changes.

**FIG. 4.**
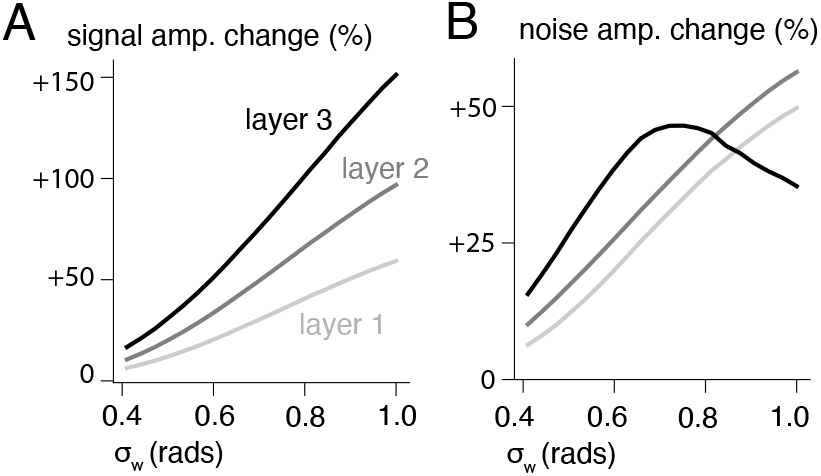
MP learning-induced changes to signal and noise. **A, B** PL-induced changes to signal (A) or noise (B) amplitude across layers for different weight selectivity. Changes are generally greater in higher layers and in networks with initial weights that are less selective (larger *σ_w_*). In both panels, *σ_s_* = 0.4, *σ_w_* = 1.0, *N* = 1000, *L* = 3.

### Impact on discrimination around untrained stimuli

MP plasticity breaks the symmetry of pre-PL representations w.r.t. *θ*, thus alters the representations of untrained stimuli. To assess how these changes affect the discrimination ability of angles around untrained values we define a normalized information gain, [*J*_*L*,post_(*θ*)−J_*L*,pre_(*θ*)]/[*J*_*L*,post_(*θ*_tr_)−*J*_*L*,pre_(*θ*_tr_)] for an untrained stimulus *θ*. Our analysis revealed a rich, non-monotonic pattern of transfer arising from MP plasticity. Consistent with experimental findings, PL increases information for stimuli similar to the trained one (“proximal transfer”, **Fig. 5**). In addition, PL also transfers to distal stimuli, where the distance between trained and test stimuli is intermediate (“distal transfer”). Importantly, PL can also decrease information for certain untrained stimuli (negative transfer), as revealed by the dips below 0 in **Fig. 5**. Finally, as expected, representations for stimuli far away from the trained one are unaffected by learning.

**FIG. 5.**
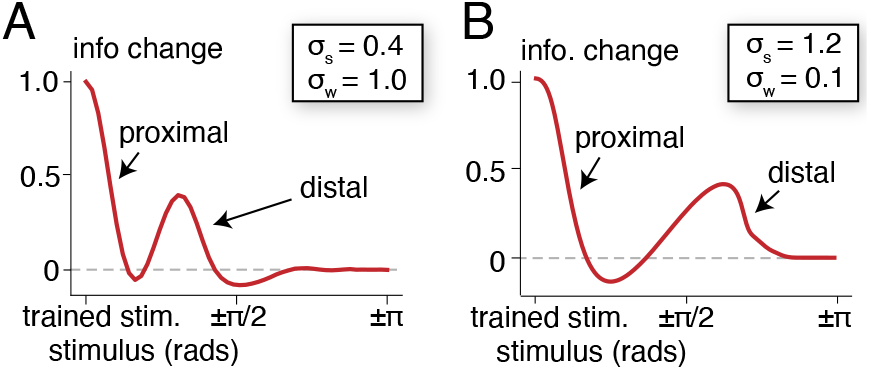
Transfer of PL to untrained stimuli. **A, B** Information changes in the last layer for different stimuli after PL, normalized by change for the trained stimulus. The change for the trained stimulus is 1 by definition. Information gain is prominent for stimuli close to the trained one (“proximal”), and those dissimilar from the trained one (“distal”). In all panels, *N* = 1000, *L* = 3, and the last layer is analyzed. For the selective-input-unselective-weights regime (**A**), *σ_s_* = 0.4, *σ_w_* = 1.0. For the unselective-input-selective-weights regime (**B**), *σ_s_* = 1.2, *σ_w_* = 0.1.

## V. MP LEARNING WITH GRADIENT DESCENT

So far, our analysis has focused on properties of MP plasticity without addressing the important question of *how* such plasticity is learned. We modeled the process of learning by studying gradient descent (GD), which has been shown to reproduce physiological features of PL in deep network models [18, 43]. We used GD to optimize Θ for a regularized loss function, defined as

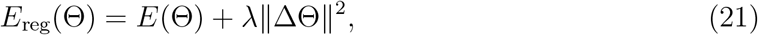

where the second term imposes a penalty on weight perturbation; strength of the penalty is controlled by the hyperparameter *λ*. We implemented GD by iterating Θ(*t* + 1) = Θ(*t*) − *η*∇_Θ(*t*)_*E*_reg_(Θ) until convergence, where *η* is the learning rate. At convergence, larger *λ* results in smaller weight perturbations but potentially suboptimal post-PL performance. To realize MP learning, *λ* should be as large as possible without making the final performance suboptimal (**Fig. 6A**). GD with such *λ* results in weight changes that are fully consistent with MP plasticity (data not shown).

**FIG. 6.**
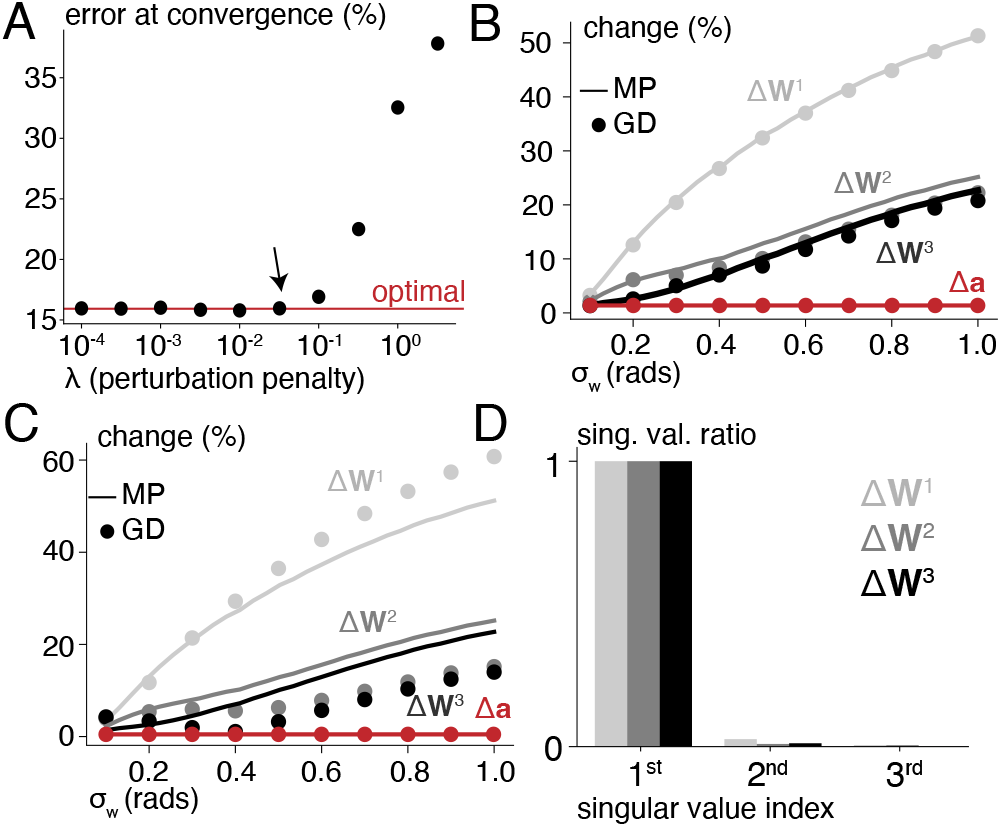
MP learning with gradient descent. **A** Discrimination error rate at convergence after regularized gradient descent with different perturbation penalty strength. Arrow: maximum *λ* with optimal performance. In all panels *σ_s_* = 0.2, *L* = 3; *σ_w_* = 0.8 except in B, D. **B** Magnitude of changes from regularized gradient descent (dots), compared against that of MP plasticity (line). **C** Same as B, but for slow GD without explicit regularization. **D** Leading singular values of of slow GD induced changes to weight matrices (normalized by the top singular value). That the first singular value is overwhelmingly large suggests that induced changes are close to rank-1.

In the deep learning literature, it has been suggested that small changes in learned parameters can also be realized through implicit regularization by using small learning rates [44, 45]. We performed GD on the unreguarlized loss function, *E*(Θ), with a small learning rate *η*. The resultant plasticity agrees reasonably well with MP plasticity (**Fig. 6C)** in terms of magnitude. It also has the same salient features as MP plasticity: changes to the readout are negligible, weight changes are very close to being rank-1 (**Fig. 6D**) and the identity of active neurons does not change over learning (S.M. V), as is the case for MP plasticity. These results point to the possibility that the slow progression of PL could be normatively explained as a mechanism to minimize perturbation during PL.

## VI. DISCUSSION

We have presented a theory of PL of fine discrimination in a deep network. The theory leverages similarity of all inputs relevant to the task, large network size and structured weight initialization to establish the effective linearity of the network input-output function during training and performance of the task. This input-output function is expressed by a processing matrix ***P*** which has been shown to be low rank, hence leading to a sub-optimal representation of the stimulus that cannot be resolved by adapting the readout weight only. We further derived the space of post-learning weights that resolve the suboptimality by fully spanning the task-relevant signal direction. Motivated by the brain’s need to strike a balance between plasticity (acquiring new skills) and stability (preventing previously learned skills to be affected) in sensory areas [46], we propose the normative minimum perturbation principle that favors a specific solution. The favored solution, which we call MP plasticity, induces physiological and behavioral changes largely consistent with current experimental findings (for a detailed comparison, see S.M. VII). It also predicts that PL improves the sensory code for some untrained stimuli while degrades the representation of others, a readily testable prediction. We discuss some prominent features of MP plasticity and their implications for neural mechanisms of PL.

First, MP plasticity predominantly modifies the lowest-level weights while leaving the readout essentially unchanged. This points to the importance of involving low-level cortical areas in PL of fine discrimination, consistent with recent numerical experiments with deep convolutional networks [43]. That the readout is unchanged critically depends on our assumption that the pre-PL readout is already optimized w.r.t. pre-learning representations, in contrast to most neural network models of learning where the initial weights are random. We argue that random initialization is not biologically plausible when considering naturalistic tasks where subjects perform well above chance with little-to-no training. Note that while synaptic plasticity is greater in lower layers, the resultant representational changes are greater in *higher* layers.

Second, MP plasticity makes rank-1 modifications to weights. While rank-1 weight changes are sufficient for optimizing neural representations for the trained task, such changes can be highly task-specific. To demonstrate this, we analyzed performance of the post-PL network on a width discrimination task where two stimuli with the same *θ* but different *σ_s_* are presented; PL does not improve this performance, despite the fact that width discrimination and angle discrimination involve the same mean stimulus (S.M. IV). Importantly, this absence of cross-task transfer reconciles the apparent inconsistency of the observed improved sensory representations by PL [1, 5, 6, 8] and the psychophysical findings that PL for one task did not transfer to another task using the same stimuli [14–16], which was interpreted as evidence that population codes for these stimuli did not improve [47]. Our results suggest that the improvement of representations does not equally benefit all tasks even if they share the same stimuli. Thus, cross-task transfer is not a reliable indicator of whether representations improve after PL.

Finally, from the perspective of signal and noise, MP plasticity improves task performance by amplifying the signal. This result is inconsistent with [19], who found that amplification is not necessary for PL. Their conclusion may be confined to the regime where performance is dominated by neural noise, not input noise as in ours. Additionally, their plasticity model differs from ours in that it assumes circularly invariant weights both before and after learning, which forces a global change of synaptic weights. In contrast, in our model, PL plasticity is localized to the neurons responding to the stimulus (if we require post-PL weights to be circularly invariant in our model, post-PL tuning curves have very unnatural multi-modal shapes. See S.M. VIII). Finally, we note that our prediction of signal amplification stems from the fact that the readout layer remains essentially unchanged under MP learning. If the readout were adapted in ways that violate the MP principle, signal amplification is not always necessary (S.M. IX).

Our current theory can be extended in several interesting directions. Our plasticity model does not include a mechanism of unsupervised learning, namely, plasticity triggered by the mere exposure to the stimulus, independent of task. Thus, including considerations for task-irrelevant plasticity, observed in some PL studies [48, 49] is an interesting topic for future work. Additionally, it would be interesting to add to our architecture recurrent connections within each layer and to consider the effect of neuronal noise on MP learning.

## ACKNOWLEDGMENTS

The authors would like to thank Andrew Saxe and Ravid Ziv for very helpful discussions. This research was partially supported by the Swartz Program in Theoretical Neuroscience at Harvard University, the Gatsby Charitable Foundation, the National Institute of Neurological Disorders and Stroke (Grant No.1U19NS104653), and the National Science Foundation (Grant No.1806818). This paper is dedicated to the memory of Mrs. Lily Safra, a great supporter of brain research.

## Supplemental Materials

**FIG. S1.**
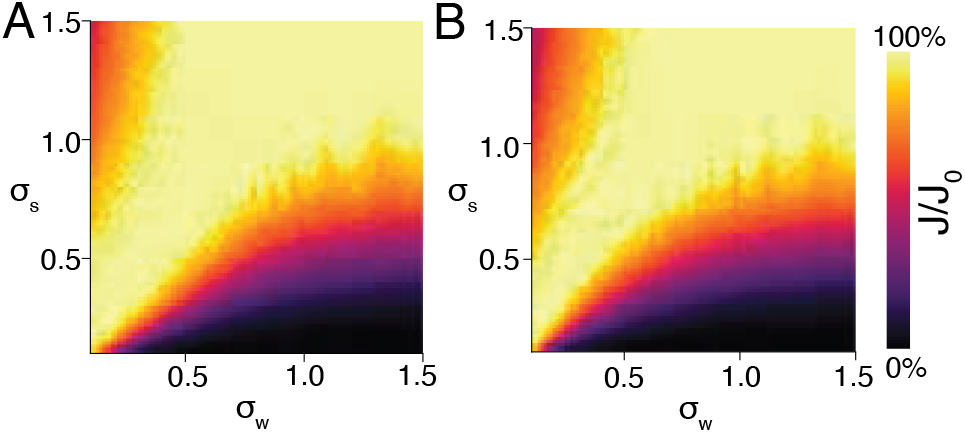
Information loss (*J/J*_0_) in untrained *L* = 2 (A) and *L* = 3 (B) networks. The *L* = 1 case is shown in Fig. 2B. The two panels share the color bar. In each case, there are two regimes corresponding to significant information loss. In all three cases, information loss is significant when the input is selective and the weights are unselective, or when the input is unselective and the weights are selective.

### I. READOUT INITIALIZATION

For brevity, we use ***W**^l^* to refer to 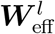 unless otherwise noted.

Under the linear approximations, the optimization problem in Eq. 7 can be solved in closed-form over ***a***. We can express the optimal ***a***_0_ as

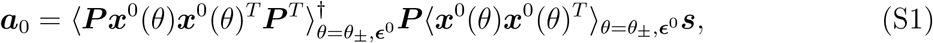

where ***P*** is defined in Eq. 13. This can be written more explicitly in term of the truncated singular value decomposition of ***P*** = ***A*****Λ*****B*** as

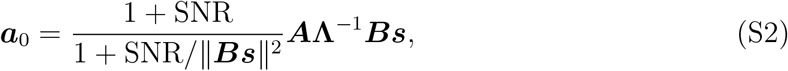

where SNR = (*δθ*)^2^||*d_θ_**f***^0^||^2^/*σ*^2^ is the input signal-to-noise ratio (SNR), which is set to 1 in simulations (see the specific values used in simulations in S.M. VI).

### II. SPACE OF SOLUTIONS

In this section we prove the sufficient and necessary condition (Eq. 16).

Formally, let **ã** = ***a**^T^**W**^L^**W***^*L*−1^…***W***^2^ ≠ **0**. We claim

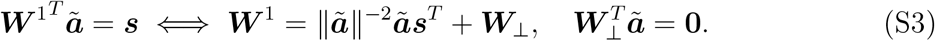

**FIG. S2.**
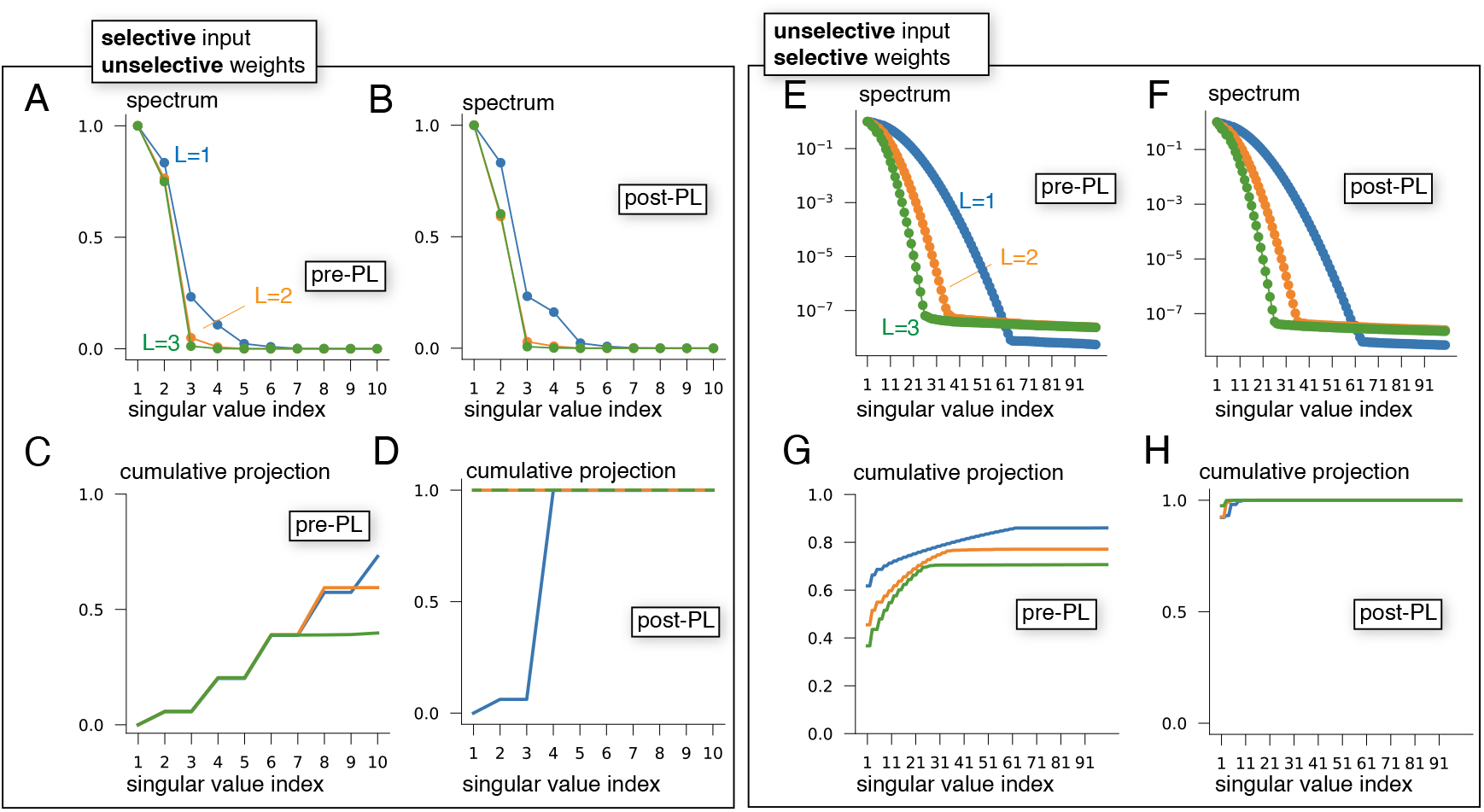
Low-rankness of weight matrices. (A) The spectrum of the product of effective weight matrices(***P***, see definition in Eq. 13) before PL. Each curve corresponds to a network of a different depth. All networks are in the selective-input-unselective-weights regime (*σ_s_* = 0.2, *σ_w_* = 0.8). ***P*** is of lower rank for deeper networks. (B) Same as (A), but for networks after PL. Rank of ***P*** for networks post-PL is approximately the same as that in pre-PL networks. (C) Sum of squared projection of the signal vector (***s***) onto the top *n* singular vectors of ***P***. (D) Same as (C), but for networks after PL. (E)(F)(G)(H) Same as (A)(B)(C)(D), respectively, but for networks in the unselective-input-selective-weights regime (*σ_s_* = 1.2, *σ_w_* = 0.1).

*Proof.* Let 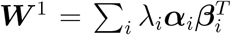 be its singular value decomposition. Then let 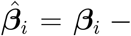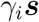 s.t. 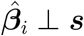 and 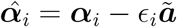 s.t. 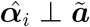. We have

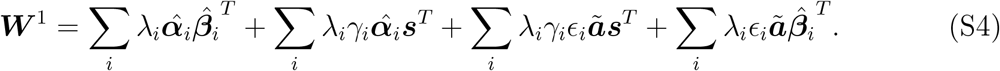

In order to satisfy 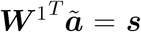, the last term must be zero. In order to satisfy 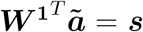, the third term must be ||***ã***||,^−2^***ãs***^T^.

Denote 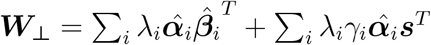,

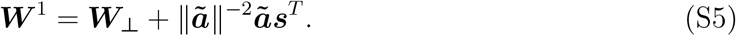

Since every component in ***W***_⊥_ has a left vector perpendicular to 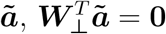.

### III. SOLUTIONS TO MINIMUM PERTURBATION (MP) LEARNING

In this section, we derive the expressions for MP plasticity, which are the solutions to the constrained optimization posed in Eq. 18, for networks of depths 1, 2, 3. These results are derived under the assumption that the same neurons are active in the pre-PL and post-PL network. The derived results are self-consistent with this assumption.

#### A. L=1

Define the Lagrangian

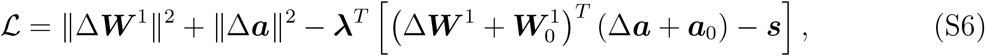

where ***λ*** is a vector of *N* Lagrange multipliers. Extremizing the Lagrangian w.r.t. Δ***W*** and Δ***a*** yields

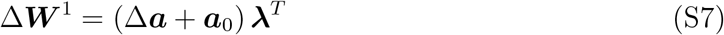

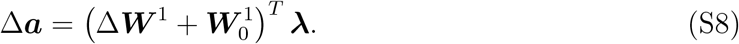

Solve for ***λ*** to get

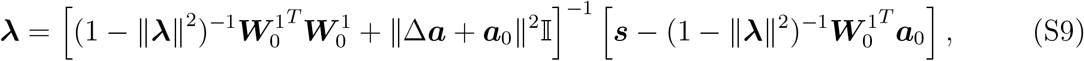

where 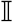 is the *N*-dimensional identity matrix. Defining scalar order parameters

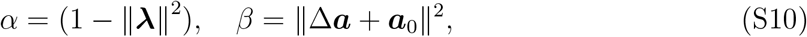

we have

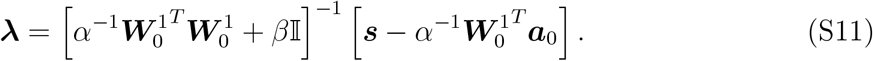

This expression can be plugged back into definitions of *α, β* to obtain two self-consistent equations for *α, β*. Values of the order parameters can then be solved numerically, yielding *α*\*, *β*\*. Plugging these back into expressions for Δ***a***, Δ***W***^1^ gives the solution.

*a. Assuming fixed **a*** If we assume a fixed ***a***, the MP Δ***W***^1^ can be given in closed form as

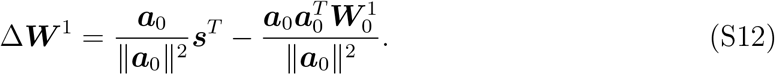

#### B. L=2

For *L* = 2, 3, applying the same analysis as above shows that changes to Δ***a*** are negligible. For simplicity, hereafter we assume Δ***a*** ≈ 0. This has the advantage of needing only 2*L* − 2 order parameters (instead of 2*L*).

Define the Lagrangian as

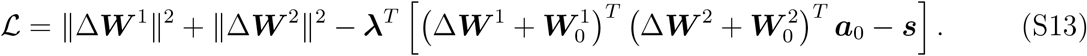

Extremizing yields

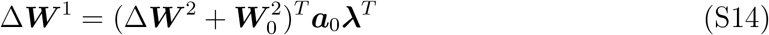

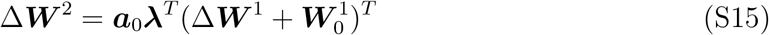

Solving for Δ***W***^1^, Δ***W***^2^ yields

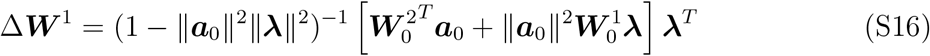

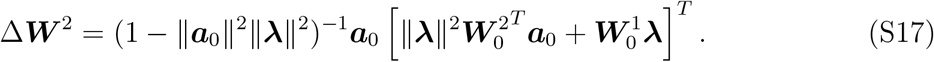

Define scalar order parameters

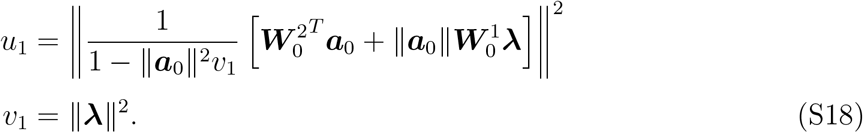

Plugging expressions for Δ***W***^1^, Δ***W***^2^ into the constraint equation and solve for ***λ*** to get

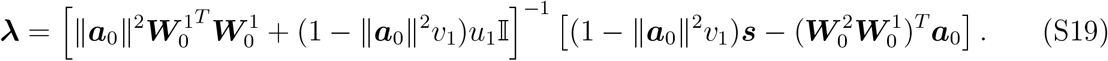

Plugging this expression back into Eqs.S18 yields two self-consistent equations of *u*_1_, *v*_1_. Other variables in these equations are all stationary in time. Therefore, one can numerically solve for *u*_1_, *v*_1_ to obtain expressions for Δ***W***^1^, Δ***W***^2^.

#### C. L=3

Setting up the Lagrangian and extremizing the variables to get

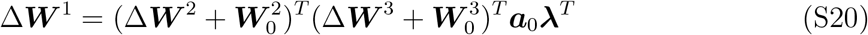

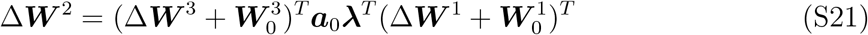

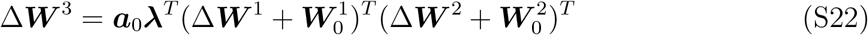

Solving these equations for Δ***W**^l^* gives (*u*_1,2_, *v*_1,2_ being order parameters defined below)

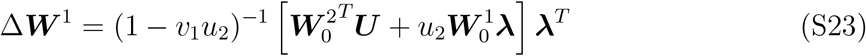

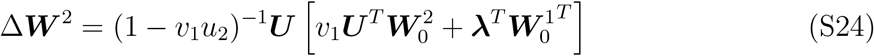

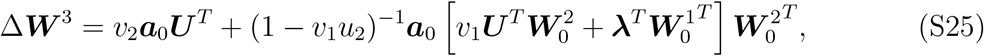

where

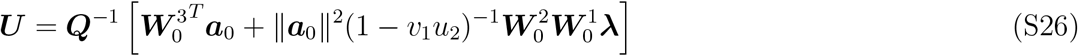

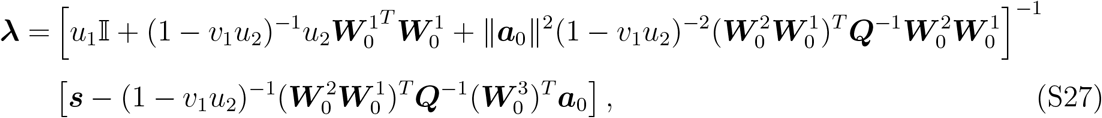

and

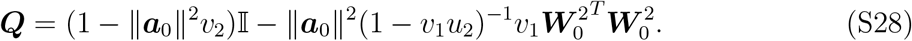

We have defined four scalar order parameters to be solved numerically.

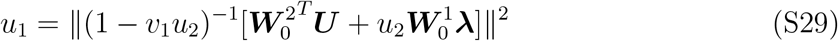

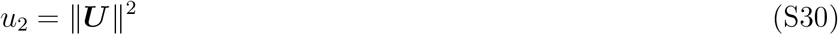

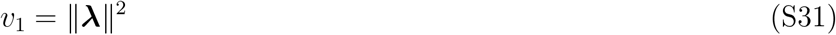

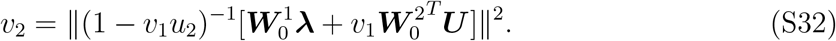

#### D. Numerical solvers of self-consistent equations

In each case discussed above, we seek to solve *k* nonlinear equations of *k* scalar variables numerically. There are many algorithms for this purpose. In general, convergence to the true solution is not guaranteed and depends on initial estimates. To obtain good initial estimates, we used a two-step procedure to solve the equations for each set of network parameters.

In the first step, we used an iterative algorithm defined in Algorithm1.

To aid convergence, we replaced all matrix inversions in the equations with pseudo-inverses (specifically, we only keep the 4 leading singular values and inverse them). These equations are referred to as “pseudo-self-consistent equations”. We first used trial-and-error to find good initial estimates for a specific set of network parameters (e.g., *σ_s_* = 0.1, *σ_w_* = 1) and ran the algorithm until convergence. We then considered another pair of parameters that are close to the previous pair (e.g., *σ_s_* = 0.125, *σ_w_* = 1), using *final estimates* for *σ_s_* = 0.1, *σ_w_* = 1 as initial estimates for the new pair. We repeated this procedure recursively to cover all network parameter regimes of interest. After this step, we obtain solutions to the pseudo-self-consistent equations.

##### Algorithm 1: Algorithm for solving self-consistent equations.

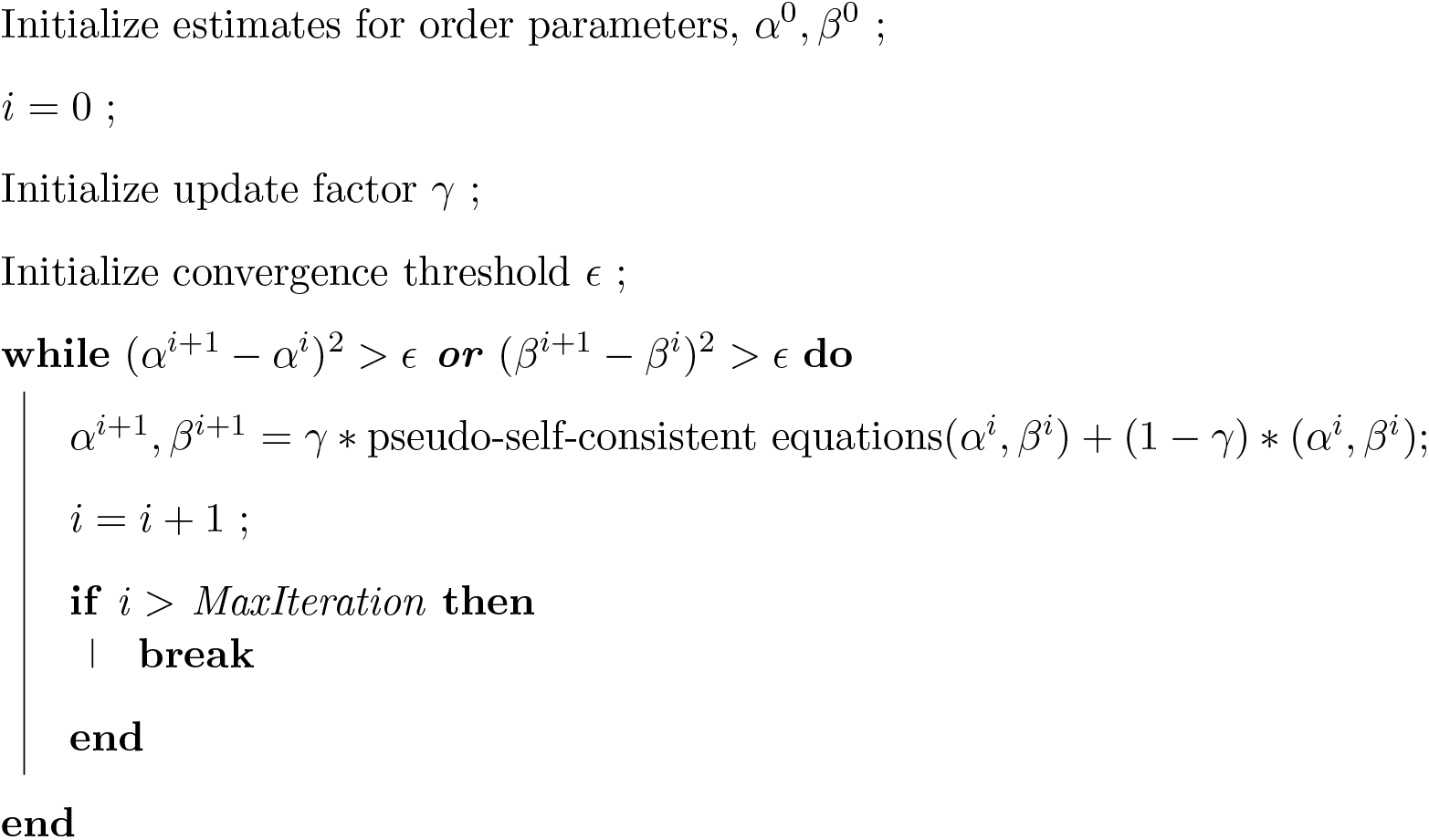

For the second step, we used `scipy.optimize.fsolve`, which implements a quasi-Newton method. For each set of network parameters, we used solutions to the pseudo-self-consistent equations as initial estimates for solutions to the true self-consistent equations. Upon convergence, we obtain solutions to the true self-consistent equations.

### IV. CROSS-TASK TRANSFER TEST

To test whether perceptual learning on the *θ* discrimination task transfers to a different task, we devised a *σ_s_* discrimination task (“width discrimination”). In this task, the network has to discriminate between two close-by values of *σ_s_*, *σ_s,_*_tr_ ± *δσ_s_* with the same *θ*. We ensure that the two tasks have the same difficulty (i.e., the optimal accuracy is the same for both tasks) by choosing values of *δσ_s_* such that 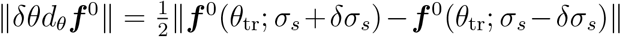.

For typical network parameters that we considered, the performance on *σ_s_* discrimination is suboptimal prior to training, leaving room for learning. Finally, we assume that the network uses two separate linear readouts from the last layer, ***a*** for *θ* discrimination and ***a***′ for *σ_s_* discrimination. Importantly, the averaged stimuli in the *σ_s_* discrimination task and the *θ* discrimination task are both ***f***^0^(*θ*_tr_).

To test for the presence of cross-task transfer, we consider the network with weights before and after MP perceptual learning and tested its width-discrimination performance with ***a***′.

Our results suggest that PL for *θ* discrimination does not transfer to *σ_s_* discrimination. This does not suggest that cross-task transfer *cannot* occur but merely provides an example where it *does not* occur despite extensive representational changes.

### V. STATIONARITY OF ACTIVE NEURONS UNDER GRADIENT DESCENT LEARNING

In this section, ***W**^l^* refers to the full weight matrix, not the effective matrix 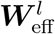. Here we show that, under GD dynamics on the unregularized loss function, ***W**^l^**f**^l^*^−1^(*θ*_tr_) is in fact stationary in time for all *l*. Our overall strategy is to show that for any *l*, ***W^l^W***^***l−1***^…***W***^***1***^***f***^***0***^, which gives a vector of synaptic inputs for neurons in the *l*th layer, is stationary in time. We first show this in networks with one hidden layer before generalizing it to deeper networks.

For brevity, denote

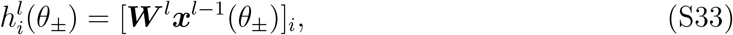

and write ***f***^0^ = ***f***^0^(*θ*_tr_). Input to the network can be written as

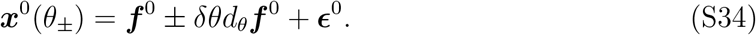

#### A. Networks with one hidden layer

In this case, output of the network is given by

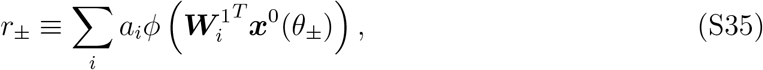

where 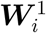 is the *i*th row of ***W***^1^. Rewrite Eq. 7 as

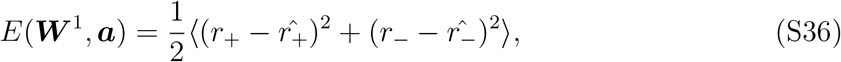

where 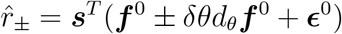 and 〈·〉 denotes average over noise.

Gradient for 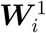 is given by

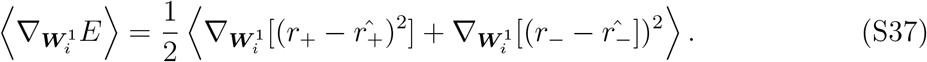

For brevity, we only examine the first term (analysis of the second term is very similar).

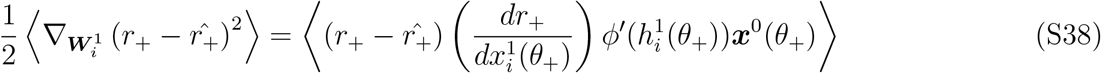

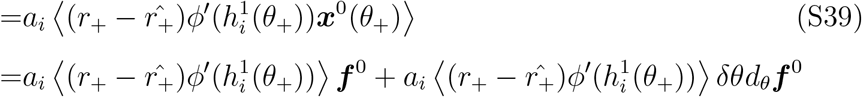

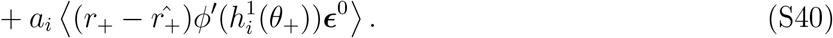

Applying Stein’s lemma to the last term yields

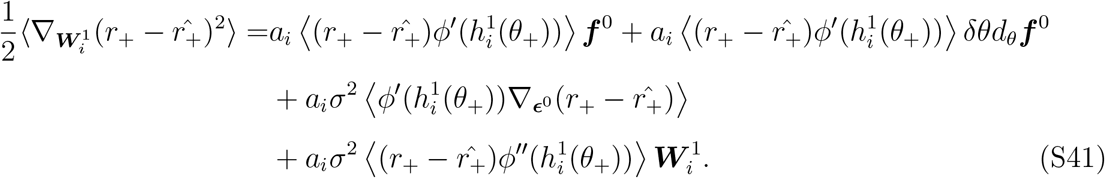

Every 〈·〉 contains the product of two random variables, which can be written as 〈*XY* 〉 = 〈*X*〉〈*Y*〉 + *cov*(*X, Y*). At large *N*, it can be verified that each is dominated by 〈*X*〉〈*Y* 〉. Eliminating covariance terms and assuming that 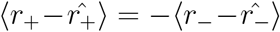 throughout training, Eq. S37 becomes

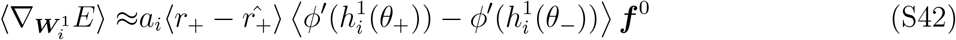

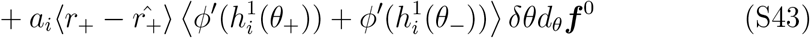

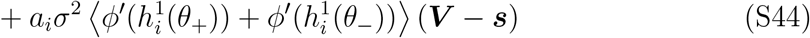

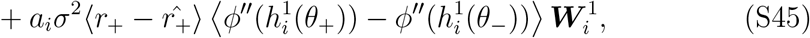

where 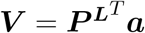. Under the linear approximations, we assume 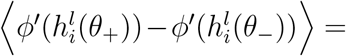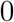. Thus,

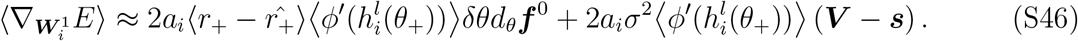

***V*** is perpendicular to ***f***^0^ because otherwise the network would be a biased discriminator. In addition, *d_θ_**f***^0^ ⟂ ***f***^0^. Thus, 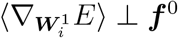 and ***W***^1^***f***^0^ is stationary in time during learning.

#### B. Generalization to deeper networks

To show that the averaged input is stationary in deeper networks, we first derive a general expression for the dynamics of ***W**^l^* during training in a deep network. Dynamics for ***W**^l^* can be derived from

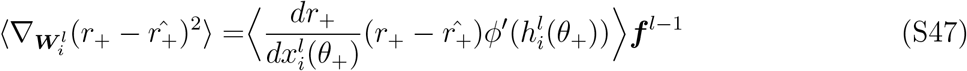

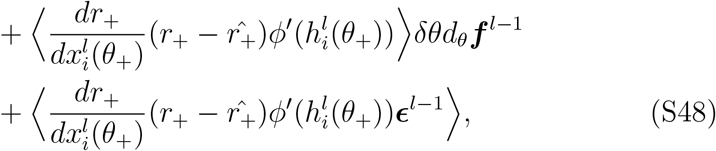

where 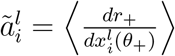 is a random variable with *O*(*N*^−1/2^) mean and fluctuation. Applying Stein’s lemma to the last term and eliminating non-leading order terms to get (let 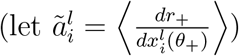)

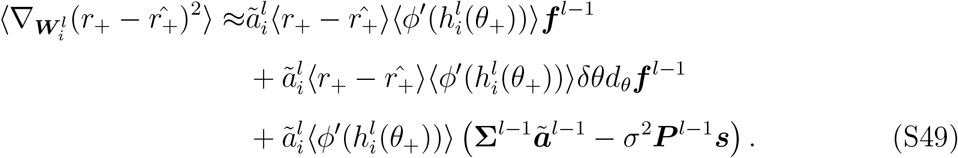

Combine 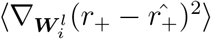 and 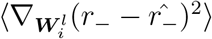 to get

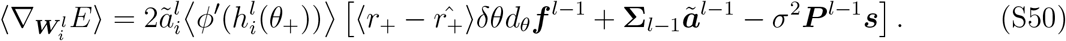

Note that under mean-field approximations, 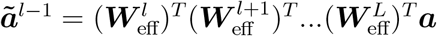. We have

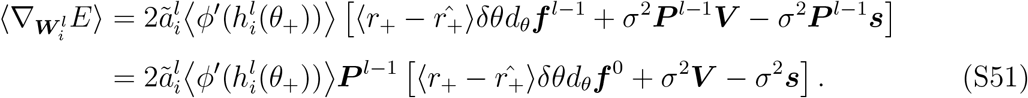

We proceed to show that every component of 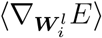 is perpendicular to ***f***^*l*−1^. First, define a notion of parity for vectors. For an *N*-dimensional vector ***v***, it is odd if for all *j*, *v*_*N*/2−*j*_ = −*v*_*N*/2+*j*_; we call it even if *v*_*N*/2−*j*_ = *v*_*N*/2+*j*_. Without loss of generality, we consider the scenario where the input neuron preferring the trained stimulus has index *N*/2. It is easy to see that *d_θ_**f***^0^ is an odd vector while ***f***^0^ is an even vector. Furthermore, any odd vector is perpendicular to any even vector.

We make the ansatz that throughout training, ***V*** is an odd vector. If ***V*** was not odd (that is, it is even or the sum of even and odd vectors), its even component would be perpendicular to the signal *d_θ_**f***^0^. This component would therefore be suboptimal because it would contribute to noise without contributing to signal. Unlike other sources of suboptimality discussed in the main text, this component can easy be removed by making ***a*** an odd vector. Since we optimize ***a*** before learning, this component does not exist at the beginning of learning.

Since pre-PL weight matrices are circulant, it is easy to verify that they have the following property: if ***v*** is an odd/even vector, then ***W**^l^**v*** and (***W**^l^*)^*T*^***v*** are also odd/even. Further, it can be verified that pre-PL effective weight matrices have the same property. We say that these matrices *preserve vector parity*. Product of matrices preserving vector parity preserves parity itself.

We now show that throughout learning, weight matrices and effective matrices preserve vector parity. Assume that at time *t*, weight matrices still preserve vector parity. Note that the gradient is a rank-1 matrix. The right vector of the gradient is an odd vector, since *d_θ_**f***^0^, ***V**, **s*** are all odd vectors, and the product matrix preserves parity. The left vector, which can be written as the element-wise product between an odd vector 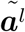 and an even vector 〈*ϕ*′(***h**^l^*(*θ*_+_))〉, is an odd vector. Therefore, for a small *τ*, this matrix at time *t* + *τ* can be written as

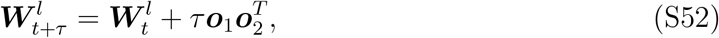

where ***o***_1,2_ are odd vectors. This new matrix will again preserve vector parity since (letting ***o*** denote any odd vector)

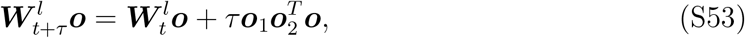

is a sum of two odd vectors and therefore still an odd vector and (letting ***e*** denote any even vector)

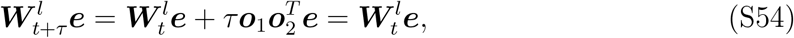

which is even. Since weight matrices preserve parity at initialization, we can show by induction that they do so throughout learning.

We now return to Eq. S51. Since weight matrices at all times preserve vector parity, the mean response in layer *l* − 1, 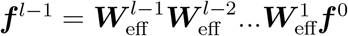 is always even.

In addition, 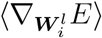 is always odd because ***P***^*l*−1^ always preserve parity. Therefore, 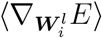 is always perpendicular to ***f***^*l*−1^(*θ*_tr_).

**TABLE S1.** Default numerical hyperparameters.

| Variable | Value | Comments |
| --- | --- | --- |
| [0.5ex] Number of neurons per layer ($N$) | 1000 | All layers have the same number. |
| Learning rate ($\eta$) | 0.001 | |
| Input noise variance ($\sigma^2$) | 0.01 | |
| $\delta\theta$ (in rads) | $\approx 0.0009$ | Adjusted so that signal-to-noise ratio in input is 1. |
| Minibatch size | 50 |  |

### VI. NUMERICAL SIMULATIONS OF GRADIENT DESCENT LEARNING

Gradient descent with minibatches was used for all weights in the network (***W**^l^* and ***a***), implemented with `pytorch`. The algorithm is terminated after the error estimated with 50, 000 examples is less or equal to the optimal level for more than 5 times. Default hyperparameters used in simulations are tabulated in Table S1. We chose relatively small *δθ* and *σ*^2^ so that the large *N* effects are already apparent with *N* = 1000, conserving computational resources. Python codes are available at https://github.com/hzshan/perceptual_learning.

### VII. DETAILED COMPARISONS WITH EXPERIMENTAL EVIDENCE

Our finding that PL is driven by improved sensory coding is consistent with the observed PL-induced changes to sensory representations in several electrophysiological experiments [1–8, 39–41] and functional imaging studies [50–53] across different model systems and tasks. Some of these studies reported representational changes that are closely related to behavioral improvements [6–8, 39], consistent with our predictions. Furthermore, we predict that for a fine discrimination task, processing between layers is approximately linear and thus all information present in an area is accessible to a linear decoder, as reported by [6]. However, our theory is inconsistent with studies that found little to no neural plasticity correlates of PL in sensory areas [2, 11, 20, 54]. Such inconsistency may arise from different task conditions and analysis methods. First, while MP synaptic changes are stronger in early layers (**Fig. 3A**), changes to neuronal tuning properties are predicted to be more prominent in higher layers (**Fig. 4**). This is consistent with experiments reporting bigger changes in monkey V4 than those in V1 and V2 following orientation discrimination PL [1–4]. This may explain why some experiments failed to find significant changes in early sensory areas [2, 54]. Second, our theory predicts that PL-induced changes to tuning curves are localized near *θ*_tr_ and may cause some tuning curves to lose their pre-PL bell shapes. Thus, studies (e.g., [2, 54]) that excluded non-bell-shaped neurons from analysis or fitted bell-shaped functions to tuning curves may fail to detect these localized changes. Third, for motion direction discrimination in moving random dot patterns [11, 20], the relevant primary sensory layer may be MT rather than neurons in V1 with smaller receptive fields. Under this interpretation, changes in the readout layer from MT should be sufficient to yield an optimal performance.

The *types* of changes to sensory representations predicted by MP learning are largely consistent with experimental observations. At a single-cell level, MP learning causes localized sharpening of tuning curves and thus amplified signal strength, as observed in experiments [1, 3, 4, 7, 39–41]. In addition, PL is predicted to decrease the number of neurons preferring *θ*_tr_, as reported in [2–4]. On a population level, MP learning decreased mean noise correlation (Fig. S4), consistent with findings from simultaneous recordings of multiple units [5–8]. We predict that mean firing rates of neurons responding to *θ*_tr_ will be unaffected by learning, consistent with [5, 11, 41, 52, 54], while some others found increased [3, 50, 51, 55–59] or decreased activation [2, 60]. While these inconsistencies may arise from differences in tasks and setups, our theory indicates that increased/decreased activation does not play a causal role in PL [6, 19].

In terms of psychophysics, our theory predicts a rich pattern of cross-stimuli transfer. While PL transfers to stimuli highly similar to *θ*_tr_ as expected, it causes performance for intermediate stimuli (“proximal”, **Fig. 5**) to drop below pre-PL levels. Indeed, some experiments report worse-than-baseline performance when subjects are tested on untrained stimuli following PL [61–63]. In addition, PL transfers to stimuli further away (“distal”) from *θ*_tr_. A more systematic examination of how observed cross-stimuli transfer depends on stimulus similarity can further test our theory. In deriving our results, we have assumed a high-precision scenario where both signal and noise are small. A sufficiently large signal (i.e., low precision) and/or noise invalidate our assumption that a fixed subset of neurons are active throughout learning as well as during responses to the trained stimuli. PL under such conditions likely involves a broader subset of neurons and weights, potentially explaining why it leads to broader cross-stimuli transfer than high-precision PL does [64].

**FIG. S3.**
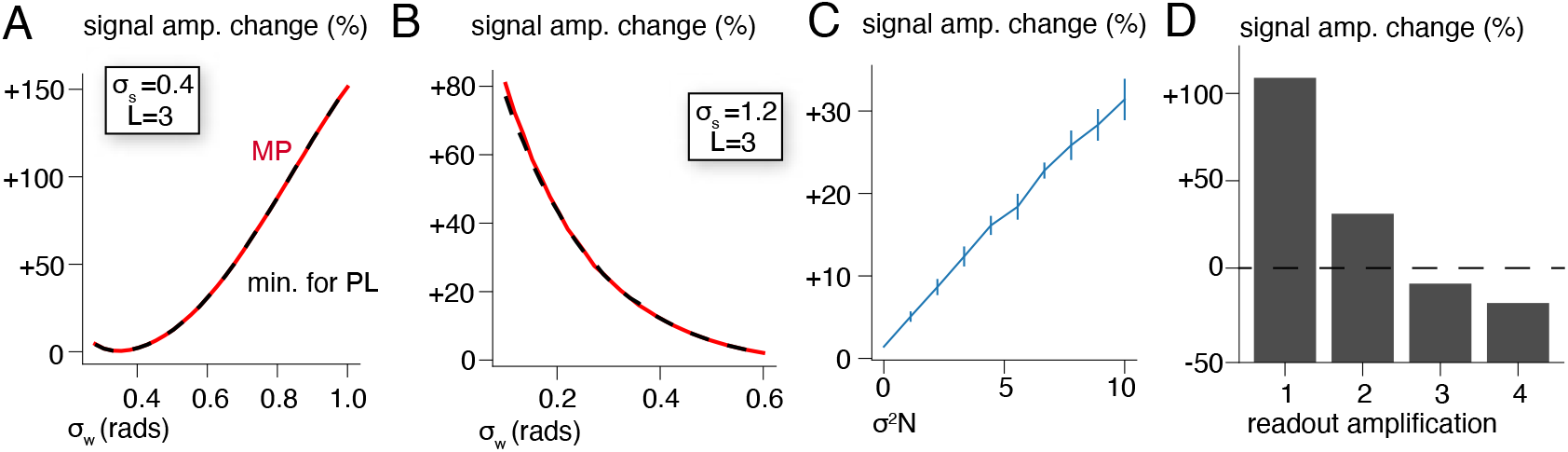
(A,B) comparison between the MP-induced signal amplification and the minimal signal amplification required for PL. (A) shows results in the selective-input-unselective-weights regime; (B) shows results in the unselective-input-selective-weights regime. (C) Average signal amplification induced by “soft” MP learning, where weights are allowed to fluctuate around the MP weights. *σ*^2^: magnitude of fluctuation. Errorbars show standard deviations over 10 independent samples. Results are taken from a 1-layer network with *σ_s_* = 0.4, *σ_w_* = 1.0. (D) Signal amplification induced by MP learning if we amplify the pre-PL readout weights. Results are taken from the last layer in a network with *σ_s_* = 0.4, *σ_w_* = 1.0, *L* = 2.

### VIII. CIRCULANT SOLUTIONS TO PERCEPTUAL LEARNING

For networks with one layer and a fixed ***a***, we seek a circulant 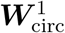 such that 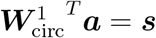. Note that all circulant matrices can be diagonalized as 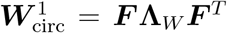, where singular vectors in ***F*** are the Fourier bases. We then use

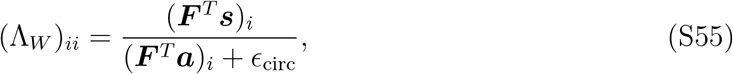

where the regularizer *ϵ*_circ_ is chosen to be as large as possible without significantly increasing 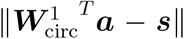. We found that such 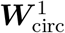, computed numerically, leads to multi-modal tuning curves, which are unrealistic results (data not shown).

### IX. IS SIGNAL AMPLIFICATION REQUIRED FOR PL?

#### A. The minimal signal amplitude in the last layer required for PL

Both MP learning and gradient-descent learning lead to negligible changes to the readout vector ***a***. Assuming the readout to be fixed, we ask whether it is possible to complete PL without increasing the signal amplitude in last-layer representations by computing the minimal signal amplitude in the last layer required for PL.

For PL, the necessary and sufficient condition in Eq. 16 translates to analogous conditions on the post PL value of the product matrix ***P**^L^*, i.e., 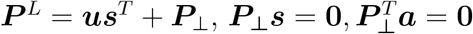. In addition, in order to minimize the loss function(Eq. 7), ***u**^T^**a*** = 1. The squared signal amplitude in the last layer is given by

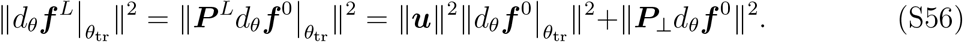

This is minimized (under the constraint that ***u**^T^**a*** = 1) by ***u*** = ||***a***||^−2^***a*** and ***P***_⊥_ = **0**. Thus, the minimal post-PL signal amplitude is

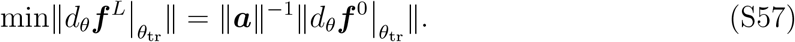

As shown in Fig. S3A,B, the minimal post-PL signal amplitude is larger than the pre-PL one across parameters. Thus, signal amplification is indeed necessary for PL, assuming a fixed ***a***. We also note that the signal amplitude after MP learning is close to the minimal level.

#### B. PL with an amplified readout vector

A reasonable extension of our model is to consider the case where the direction of the readout vector stays the same but its amplitude is increased, namely (***a***_post_ = (1 + *c*)***a***_pre_). We computed MP changes to weight matrices for several values of *c* (for networks with *L* = 2) and find that if *c* is sufficiently large, MP learning may even lead to a *decreased* signal amplitude (Fig. S3C).

#### C. Learning under a soft MP constraint

Compared to a non-MP post-PL solution to PL, does MP learning lead to smaller signal amplitudes? We consider a “soft” MP constraint where changes to the weights fluctuate in the space of solutions around MP changes. Concretely, for networks with one-layer and a fixed ***a***, we sample solution ***W***^1^ with where 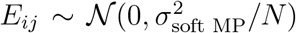 and 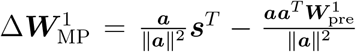 is the MP changes to ***W***^1^ (derived under S.M. III).

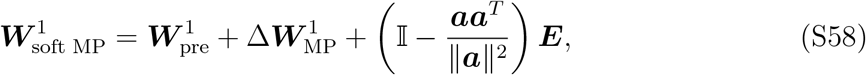

**FIG. S4.**
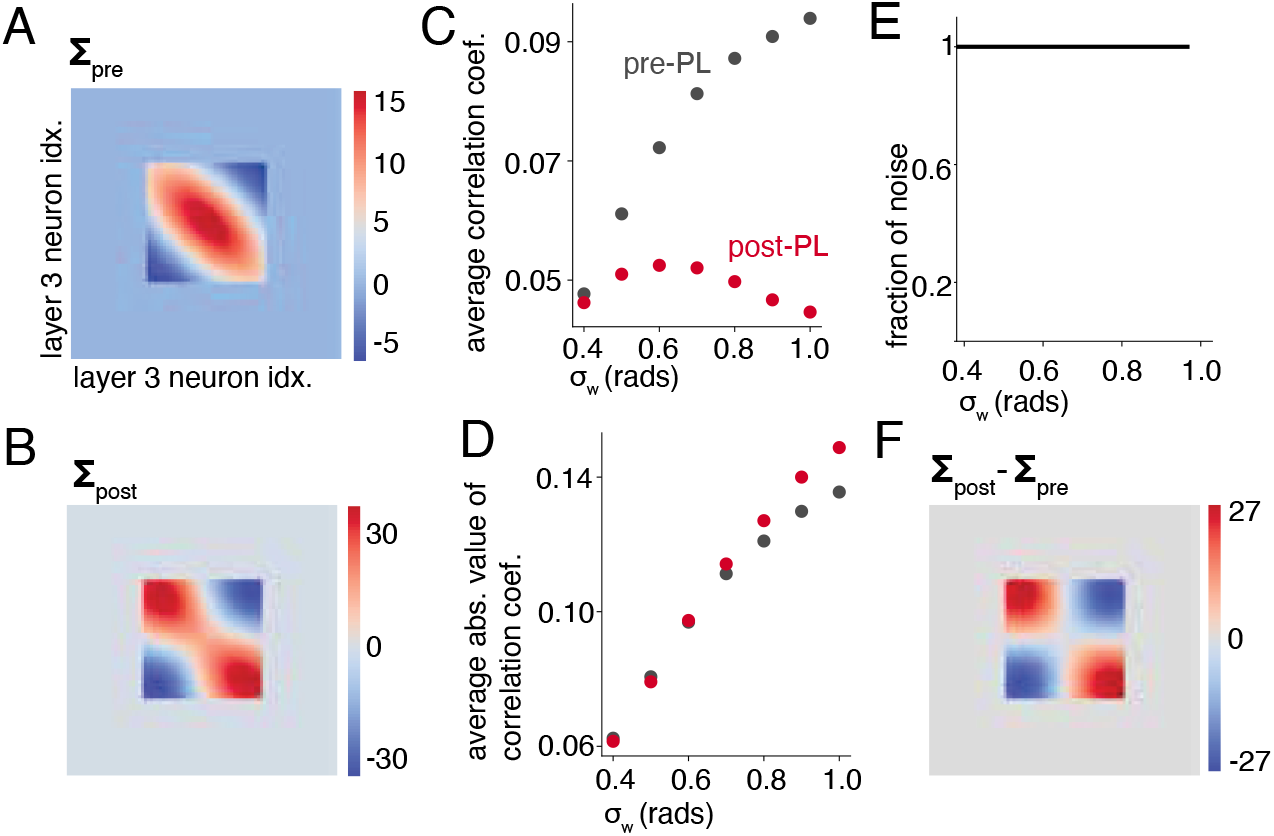
Structure of noise correlation in layer 3 of a *L* = 3 network when *θ*_tr_ is presented. (A) Normalized* pre-PL noise covariance matrix. (B) Post-PL covariance. (C) Averaged pair-wise Pearson’s correlation coefficients before (red) and after PL (black), for different *σ_w_*. (D) Same as C, averaged absolute values of coefficients are shown. (E) Fraction of PL-induced changes to covariance (**Σ**_post_ − **Σ**_pre_) that project on the signal direction. (F) Visualization of PL-induced changes to covariance. *Covariance matrices are multiplied by *N/σ*^2^||***f***^0^||^2^||***f***^3^||^−2^, where ***f***^0^, ***f***^3^ are noise-averaged population response vectors in the input layer and layer 3, respectively. This scaling makes each element of the matrix *O*(1) and adjusts for different activity levels in different layers.

Every sampled ***W***^1^ solves PL. We computed the post-PL signal amplitude for various 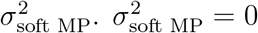 is the MP changes (Fig. S3D). We found that MP learning does indeed lead to a smaller signal amplitude than the average “soft” MP learning.

**FIG. S5.**
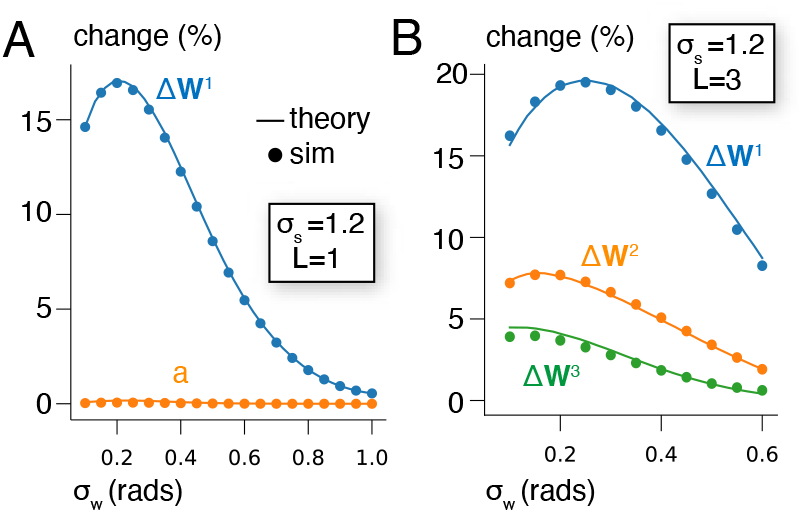
Comparing magnitude of MP learning changes (“theory”) and changes induced by slow gradient descent (“sim”) in the unselective-input-selective-weights regime. (A) L=1 network. (B) L=3 network. Learning rate is 10^−3^.

